# Automated improvement of stickleback reference genome assemblies with Lep-Anchor software

**DOI:** 10.1101/2020.08.18.255596

**Authors:** Mikko Kivikoski, Pasi Rastas, Ari Löytynoja, Juha Merilä

**Author notes:** Correspondence Mikko Kivikoski, Faculty of Biological and Environmental Sciences, FI-00014 University of Helsinki, Finland., Ari Löytynoja, Institute of, Biotechnology, FI-00014 University of Helsinki, Finland. These authors contributed equally to this work (Mikko Kivikoski, Pasi Rastas).

## Abstract

We describe an integrative approach to improve contiguity and haploidy of a reference genome assembly and demonstrate its impact with practical examples. With two novel features of Lep-Anchor software and a combination of dense linkage maps, overlap detection and bridging long reads we generated an improved assembly of the nine-spined stickleback (*Pungitius pungitius*) reference genome. We were able to remove a significant number of haplotypic contigs, detect more genetic variation and improve the contiguity of the genome, especially that of X chromosome. However, improved scaffolding cannot correct for mosaicism of erroneously assembled contigs, demonstrated by a de novo assembly of a 1.7 Mbp inversion. Qualitatively similar gains were obtained with the genome of three-spined stickleback (*Gasterosteus aculeatus*). Since the utility of genome-wide sequencing data in biological research depends heavily on the quality of the reference genome, the improved and fully automated approach described here should be helpful in refining reference genome assemblies.

## INTRODUCTION

Great deal of present-day research in biology is based on genomic data that are processed and analyzed in the context of a linear reference genome. Typical examples of this are whole-genome sequencing studies where sequencing reads are mapped to the reference genome and the characteristics of interest are derived from local dissimilarities and statistics based on the alignments (Korneliussen, Albrechtsen, & Nielsen 2014; Schraiber & Akey 2015). Reliability of those characteristics and the conclusions drawn from them depend not only on the quality of the sequencing data but also on the quality of the reference genome. Assembling and evaluating the quality of reference genomes is not easy (Baker 2012; Church et al. 2011; Meltz Steinberg et al. 2017; Rice & Green 2019). The profound problem is that the physical connectivity is lost during sequencing and recovering that in the assembly stage is notoriously difficult. To this end, high-quality linkage maps are valuable and allow inferring the physical order and orientation of the assembled contigs (Pengelly & Collins 2019; Rastas 2020; Stemple 2013).

Although a linear reference genome is ill-suited for describing many structural variations, most genome analysis methods assume the reference genome to contain each genomic region only once. The continuous development of the human reference genome (Schneider et al. 2017; Sherman & Salzberg 2020) has shown that creating a linear haploid reference genome for a diploid species is a non-trivial task. Reaching this ideal can be especially challenging in organisms where the genetic variation cannot be reduced in controlled inbreeding designs, and most reference genomes are likely based on reference individuals carrying long alternative haplotypes (Chin et al. 2016; Howe et al. 2013; Stemple 2013). Presence of homologous haplotypes, that is, differing copies of the same genomic region inherited from the two parents, is against the assumptions of the linear reference genome and affects for instance the read mapping. If reads from distinct haplotypes map to different copies of the same region, single nucleotide variants (SNPs) separating the haplotypes cannot be detected and variation is underestimated. This affects various statistics in population genomics, and may lead to wrong conclusions in many different contexts, including estimation of substitution rate (Kong et al. 2012), inbreeding (Ceballos, Joshi, Clark, Ramsay, & Wilson 2018) or population history (Roux et al. 2016).

Lep-Anchor software (Rastas 2020) can improve assembly and scaffolding of even high-quality reference genomes with joint use of linkage map based genome anchoring, pairwise contig alignment and long-read sequencing data. Performance and utility of Lep-Anchor were demonstrated in its original publication (Rastas 2020) with empirical and simulated data sets and gains in assembly quality were reported even with relatively small data sets. Here, we have a closer look on the actual changes and assess their impact on typical genome analyses. Starting from an existing high-quality contig assembly, original PacBio reads and ultra-dense linkage maps for the nine-spined stickleback (*Pungitius pungitius*), we were able to generate a significantly improved reference genome (ver. 7) using largely automated methods. When evaluating the differences to the published version of the reference genome (ver. 6; Varadharajan et al. 2019) we detected haplotypes in three contexts. First, some haplotypes were originally assembled as separate contigs leading to false duplication of a region in the assembly. Second, haplotypes were assembled to the ends of subsequent contigs and occurred as duplicates on both sides of a contig gap. Third, haplotypic regions, exemplified by an inversion in LG19, were assembled as mosaics of the two haplotypes. Using the novel features of Lep-Anchor, we could automatically remove a large proportion of the first two types of haplotypes while the correction of haplotypes of the last category was possible but demanded manual effort. Recognition and removal of haplotypes shortens the nine-spined stickleback reference genome and increases heterozygosity of the reference individual while the contig re-scaffolding enabled the identification of the centromere in all linkage groups. To demonstrate that this approach works for contig assemblies in general, we reassembled the latest published reference genome of the three-spined stickleback (*Gasterosteus aculeatus*; Peichel, Sullivan, Liachko, & White 2017) using one new linkage map and publicly available 10X Genomics linked read sequencing data (Berner et al. 2019).

## 1 MATERIALS AND METHODS

### Nine-spined stickleback reference genome refinement

The starting point for this reference was the contig assembly and the genomic DNA sequence data from Varadharajan et al. (2019). In short, the ver. 6 genome by Varadharajan et al. (2019) was based on *de novo* assembly of long PacBio reads, polishing with short reads and anchoring with linkage maps. The contig assembly was refined in two places: (1) the mitochondrial genome was reassembled from the short-read Illumina data of the reference individual using the program MEGAHIT (ver. 1.2.9; D. Li, Liu, Luo, Sadakane, & Lam 2015), and (2) a large inversion in LG19 was characterized and the region was reassembled using the combination of programs Falcon Unzip (ver. 0.4.0; Chin et al. 2016), Trio Binning (prerelease version; Koren et al. 2018), Canu (ver. 1.6; Koren et al. 2017) and Pilon (ver. 1.22; Walker et al. 2014), all run with their default parameters. The details of these steps are provided in the Supplementary methods.

A new ultra-high density linkage map was reconstructed based on crosses of wild-caught marine nine-spined sticklebacks from Helsinki, Finland (60*°* 13’N, 25*°* 11’E). 99 F_1_-generation families were generated at the University of Helsinki fish facility through artificial fertilizations (Rastas, Calboli, Guo, Shikano, & Merilä 2016). Half-sib families were formed by mating one female to two different males, thinning the families to 25 offspring per family. The larvae were mass-reared in two large aquaria and their family identity was later identified from the genotype data. The parental fish were whole-genome sequenced (WGS; Illumina Hiseq platforms, BGI Hong Kong) at 5–10x sequencing coverage and the offspring were genotyped using the DarTseq technology (Diversity Arrays Technology, Pty Ltd, Australia). The fastq files were mapped to the contig assembly using BWA-MEM (ver. 0.7.15; H. Li 2013) and SAMtools (ver. 1.9; H. Li et al. 2009). The genotype likelihoods were called and the linkage mapping and the pedigree construction were conducted using Lep-MAP3 (Rastas 2017). The details of the linkage map reconstruction are provided in the Supplementary methods.

The resulting contig-assembly was anchored using Lep-Anchor (Rastas 2020) following the standard pipeline (https://sourceforge.net/p/lep-anchor/wiki/Home) with default parameters (exception: minQuality=1 for Map2Bed to assign more contigs into chromosomes). For the anchoring, we (1) utilized three original linkage maps (Varadharajan et al. 2019) and the newly reconstructed ultra-high density linkage map concordant with the existing maps; (2) generated contig-contig alignments by running the two first steps of HaploMerger2 (Huang, Kang, & Xu 2017); and (3) incorporated the raw PacBio reads by aligning them to the contig assembly with minimap2 (ver. 2.17; H. Li 2018). Full computer code for reproducing these analyses and instructions for automated improvement of any reference genome assemblies are available at https://github.com/mikkokivikoski/NSP_V7.

### Contig classification and centromere annotation

In ver. 7, 1644 of the total 2487 contigs were not assigned to any of the 21 linkage groups (Table 1). We classified the contigs by analyzing their sequencing depth (coverage) and repeat content. Illumina and PacBio data (subreads) for the reference individual and for a pool of four female individuals from the same Pyöreälampi pond (Illumina only, see Supplementary methods for the details) were mapped and analyzed using BWA-MEM and minimap2, respectively, and SAMtools. The coverage analysis was carried out using Lep-Anchor’s novel modules CoverageAnalyser and CoverageHMM. Using CoverageAnalyser and a simple mixture model, sequencing depth histogram was classified to (about) zero, half, normal or high: Half and normal depths were modelled using two normal distributions and the zero and high depth as a zeta distribution (coverage + 1 *∼* Zeta, the same distribution was used for both, zero and high). Then CoverageHMM and a four-state hidden Markov model (HMM) were used to classify each genomic position to four states: zero, half, normal or high. The emission probabilities of the HMM were taken from the mixture model (CoverageAnalyser) and maximum likelihood transition probabilities along the physical (contig) coordinates were learned using the Baum-Welch expectation-maximization algorithm (Baum et al. 1972).

**TABLE 1.** Summary of the differences between the two nine-spined stickleback genome assemblies

| Feature | ver. 6 | ver. 7 | %Change |
| --- | --- | --- | --- |
| N50 contig size (bp) | 1,202,809 | 2,794,615 | +132.34 |
| Total length of the assembly (bp)<sup>†</sup> | 521,233,387 | 466,582,808 | -10.48 |
| Total length of the 21 linkage groups (bp) | 444,482,085 | 439,721,235 | -1.08 |
| LG12 length (bp) | 40,899,740 | 33,585,825 | -17.88 |
| Contigs in linkage groups (contig chains)<sup>‡</sup> | 686 (NA) | 843 (362) | +22.89 (NA) |
| Contigs in LG12 | 244 | 150 | -38.52 |
| Contigs not assigned to linkage groups (length) | 4,616 (76,734,720 bp) | 1,644 (27,251,636 bp) | -64.38 (-64.49) |
| Contigs not in linkage groups of other assembly (LG12) | 117 (109) | 274 (15) |  |
| Contigs with known orientation | Not assessed in ver. 6 | 763 (427,086,963 bp) |  |
| Complete BUSCOs | 3572 (98.2%) | 3573 (98.2%) | +0.03 |
| Complete single-copy BUSCOs | 3438 (94.5%) | 3529 (97.0%) | +2.65 |
| Complete duplicated BUSCOs | 134 (3.7%) | 44 (1.2%) | -67.16 |
| Fragmented BUSCOs | 18 (0.5%) | 16 (0.4%) | -11.11 |
| Missing BUSCOs<sup>§</sup> | 50 (1.3%) | 51 (1.4%) | +2.00 |
| Total BUSCO groups searched | 3640 | 3640 | 0 |

<sup>†</sup>Includes 21 linkage groups with gaps, unassigned contigs and mitochondrial sequence. <sup>‡</sup>Contig chain refers to group of contigs joined without gap in ver. 7. In ver. 6, all contigs had a gap in between. <sup>§</sup>See Suppl. Table 3

Repetitive regions were identified with RepeatMasker (ver. open-4.0.5; Smit *et al*. 2013–2015 http://www.repeatmasker.org) by using the species specific repeat libraries by Varadharajan (2019). Contigs with *>*20% repeat content were classified as repetitive contigs (Fig. 1a). The centromere-associated repeat sequence characterized by Varadharajan (2019) was aligned against each unassigned contig with blastn (BLAST+ applications version 2.2.31+; Altschul, Gish, Miller, Myers, & Lipman 1990; Camacho et al. 2009). All contigs with at least one hit with e-value *<* 10^*−*5^ were classified as putative centromeric contigs.

**FIGURE 1.**
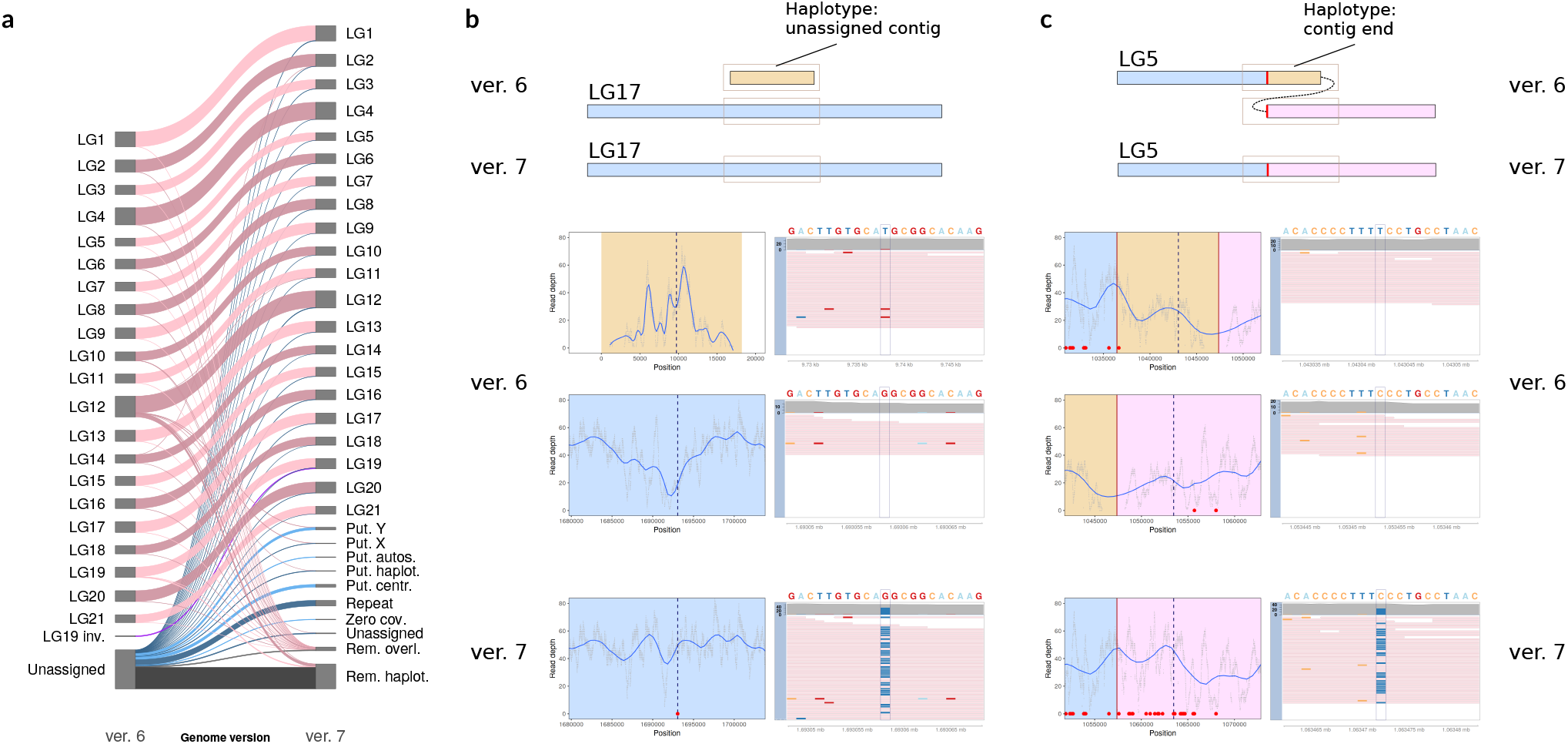
Summary of the changes between ver. 6 and ver. 7 of the nine-spined stickleback reference genome and examples of removed haplotypes. (a) Diagonal lines indicate changes in contig placement between different linkage groups (LGs) with band widths proportional to the length of the contigs with the corresponding change. Unassigned contigs in ver. 7 were grouped into putative classes according to their sequencing coverage and repeat content (see methods). (b, c) A schematic illustration of regions in the two assemblies is shown on top and the data for the highlighted areas (boxes) in the panels below. On the left, blue curves show the smoothed read depth and the dashed lines indicate a SNP position, boxed in the right panel. On the right, the reference sequence is shown on top and the pink bars indicate mapped reads, mismatches shown with matching colors. (b) A short, unassigned contig (orange) was identified as a haplotype within a contig (blue) in LG17. After its removal (ver. 7, bottom), the read depth is more even and a new SNP (red dot) is identified. (c) A region (orange) was duplicated in the ends of neighboring contigs (blue, pink) in LG5. After its removal (ver. 7, bottom; cut site in red), the read depth is more even and several new SNPs are identified.

Alignments of centromere-associated repeat sequence were used to determine the centromere positions (Suppl. Table 1, Suppl. Fig. 1). Within each linkage group, Blast alignments with e-value *<* 10^*−*5^ were assigned to three groups with k-mean clustering according to their position. Clusters with less than 10% of the total number of hits were discarded as outliers, and the centromeric region was defined to span the remaining hits. Analyses were conducted and the results visualized with R (ver. 3.4.4; R Core Team 2018 https://www.R-project.org/) using packages ggplot2 (ver. 3.0.0; Wickham 2016) and ggforce (ver. 0.3.1; Pedersen 2019 https://CRAN.R-project.org/package=ggforce).

### Content of LG12 sex chromosome and LG19 inversion

Based on the female and male sequencing coverage, the sex-chromosome part (1–25 Mpb) of the ver. 6 LG12 appeared to contain contigs derived from X and Y chromosomes. We aimed to make LG12 haploid and purely X, and to identify differentiated Y-origin haplotypes (Table 1). To investigate the new assembly of LG12 we joint-called variable sites in a pool of the reference individual and four females using GATK4 (ver. 4.0.1.2; McKenna et al. 2010), and defined a HMM based on the frequency of homozygous reference and variant alleles in females. We assumed that females are homozygous for the reference allele in regions representing X and homozygous for the variant allele in regions representing Y. The emitted statistic was 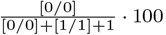, where [0/0] and [1/1] are the number of loci where an individual is homozygous for reference or variant allele, respectively. The statistic was calculated in 50 kb windows and rounded to the closest integer. Low and high values of the statistic indicate X and Y chromosomes, respectively, whereas values of around 50 indicate fine-scale mosaicism of X and Y. The analysis was carried out in the sex-chromosome region of the ver. 7 LG12 (1–16.9 Mbp) with R package HMM (ver. 1.0; Scientific Software Development, Himmelmann 2010 https://CRAN.R-project.org/package=HMM).

The two alleles for the LG19 inversion were *de novo* assembled using the long-read data from the reference individual and short-read data from related individuals homozygous for the different copies (see Supplementary methods for the details). Alternative versions of the genome were created by inserting the newly assembled alleles into the reference sequence. Individuals homozygous for the a and b alleles were mapped to different versions of LG19 with BWA-MEM and SAMtools. Variants were called with bcftools mpileup (ver. 1.9; H. Li 2011) and single nucleotide variants with quality score *≥* 5 were retained. Frequencies of sites with homozygous and heterozygous variant alleles were calculated in 100 kb windows with Bedtools software (ver. 2.27.1; Quinlan & Hall 2010).

Another HMM was defined to identify potential other inversion haplotypes. We anticipated that a dense mosaic of haplotypes in the reference genome results in variation between homozygous reference and variant alleles in an individual homozygous for one haplotype. Therefore, emitted statistic was defined 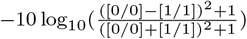 as The where [0/0] and [1/1] are the number of loci where an individual is homozygous for reference or variant allele, respectively. The statistic was estimated in 50kb windows and rounded to the closest integer; values above 40 were truncated to 40. Small values (e.g. high proportion of both homozygous genotypes) indicated inversion region. The HMM was applied to four female individuals and all 21 linkage groups.

### Quality assessment with variant and synteny analyses

To compare the nine-spined ver. 6 and ver. 7 references, we called autosomal SNPs of the reference individual (FIN-PYO-0). Reads were mapped to both references using BWA-MEM and variants were called with bcftools mpileup. SNPs were pruned with stringent criteria: SNPs within repetitive or unmappable regions, within 20 bp of an indel, of low quality (< 20) or with low (< 30) or high (> 70) depth were discarded. Unmappable regions were determined using the approach of Li (http://lh3lh3.users.sourceforge.net/snpable.shtml) and converted to bed format using a script by Schiffels (https://github.com/stschiff/msmc-tools). SNPs found using ver. 6 were grouped into three categories: (1) found in autosomal linkage groups of ver. 7, (2) locus removed from autosomal linkage groups of ver. 7, or (3) not called with ver. 7. SNPs called using ver. 7 were grouped similarly but there were two additional groups for SNPs in regions where haplotype copy was removed (Table 2).

**TABLE 2.**
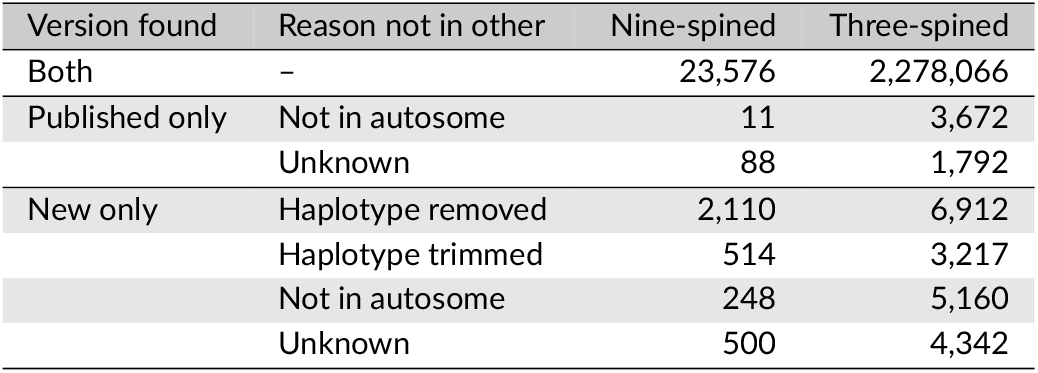
Number of autosomal SNPs of detected by mapping short read data against the published and the new assemblies. SNPs may be missing because the region is involved in haplotype removal or is excluded from the autosomes. “Unknown” indicates SNPs identified in regions with no contig changes or removed haplotypes.

The quality of ver. 6 and ver. 7 were also assessed by comparing their synteny with the three-spined stickleback genome (Peichel et al. 2017). Based on the previously reported large-scale synteny to the three-spined stickleback genome (Varadharajan et al. (2019); see also Guo, Chain, Bornberg-Bauer, Leder, and Merilä (2013); Rastas et al. (2016)), the homologous linkage groups of nine- and three-spined sticklebacks were aligned with minimap2 software. Previous studies (Rastas et al. 2016; Shikano, Laine, Herczeg, Vilkki, & Merilä 2013) have shown that LG12 is a fusion chromosome, and it was aligned against the three-spined stickleback linkage groups 7 (1–14 Mbp) and 12. Alignment fragments with less than 5000 matching base pairs were discarded and the syntenies of the two assemblies with the three-spined stickleback genome were compared by counting the number of changes in orientation of consecutive fragments (Fig. 2b). BUSCO completeness of ver. 6 was reported to be very high, containing 97.1% of tested genes as complete BUSCOs (see Table 1 in Varadharajan et al. (2019)). Here, we carried out the same analysis for both genome versions using BUSCO ver. 5.0.0 (Seppey, Manni, & Zdobnov 2019). The command used was ‘docker run −u $ID −v $PATH:busco_wd ezlabgvabusco:v5.0.0_cv1 busco −m genome −i reference.fasta −o result_busco_reference –auto-lineage-euk’. Contig classification, variant analysis, synteny comparisons and other downstream analyses of the genome assembly, were executed using Anduril 2 workflow platform (Cervera et al. 2019).

**FIGURE 2.**
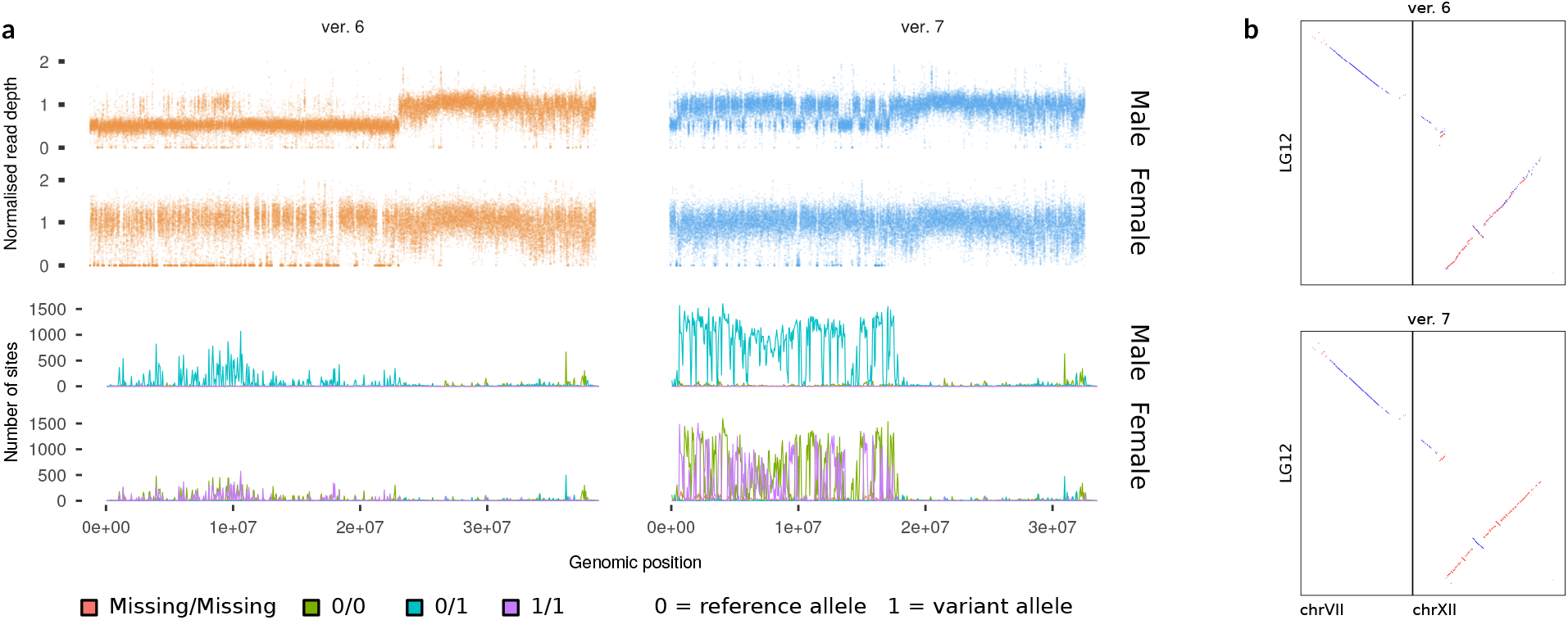
Improvements in LG12 sex chromosome. (a) Normalised read depth (top) of the male reference individual (50x coverage) and a female (10x, FIN-PYO-20) is closer to the expected (one) in ver. 7 (right) and fewer regions show zero depth. Ver. 7 has more segregating sites (bottom) and especially sites where the reference individual is heterozygous (turquoise) and the female is homozygous (green, purple). Number of sites is calculated in 100 kb windows. (b) The synteny of the nine-spined stickleback LG12 with the three-spined stickleback genome (x axis) is more contiguous in ver. 7, and there are fewer changes in contig order. Red and blue colors indicate forward and reverse alignments, respectively.

### Three-spined stickleback reference genome refinement

We also tested the performance of Lep-Anchor with the three-spined stickleback genome assembly (Peichel et al. 2017). First, a linkage map was constructed with Lep-MAP3 based on the data set of 517 F_1_-offspring from 60 families (30 males, each crossed with two females) described by Pritchard et al. (2017). The parents were wild caught from the Baltic Sea and artificially crossed (see Leder et al. (2014) and Pritchard et al. (2017) for more details). The linkage map reconstruction differed from that of the nine-spined stickleback in two places: the pedigree was obtained from Pritchard et al. (2017) and, in Separate-Chromosomes2, lodLimit was set to 25 to obtain 21 linkage groups.

The original scaffolded genome was partitioned into (about 16,000) underlying contigs by cutting it at long runs of N’s. An artificial map was made to contain one marker per contig, listing contigs in the scaffold order within each of the 21 linkage groups. To allow deviations from the contig order of Peichel et al. (2017), the marker for the i:th contig was given a map interval of [i, i+9]. Finally, an artificial alignment file (paf format) was constructed with alignments for each adjacent contig in the scaffolds. As for the nine-spined stickleback, we then run the Lep-Anchor pipeline using the linkage map produced with Lep-MAP3 and the artificial map and alignment files. In the lack of long-read data, we incorporated a scaffold level 10X Genomics genome assembly (Boot Lake population, Vancouver Island, Canada; Berner et al. 2019) into the input data. The 10X assembly and the three-spined stickleback contigs were aligned with minimap2 and included as two copies to Lep-Anchor to increase its weight in the optimisation score.

Lacking the short-read data for the reference individual, we called SNPs for a male three-spined stickleback from Paxton Lake benthic population, Canada (Samuk et al. 2017). The Illumina WGS data for the sample SRR5626529 were downloaded from European Nucleotide Archive (ENA) and mapped with BWA-MEM to the published three-spined stickleback genome and to the genome assembled here. SNPs were called with bcftools mpileup as in the nine-spined stickleback (see above). As the mean sequencing coverage of the sample as 15x, only SNPs with depth between 7 and 23 were retained.

## RESULTS

We used Lep-Anchor software and information from linkage map anchoring, pairwise contig alignment and long-read bridging to reassemble the nine-spined stickleback genome. Linkage map anchoring allowed assigning 274 previously unassigned contigs to the linkage groups (LGs) and pairwise contig alignments revealed 10% of the previous assembly as haplotypes (Fig. 1a). Of the 843 contigs in linkage groups, Lep-Anchor could assess 763 to be scaffolded in correct orientation. Removal of haplotypes and linking of adjacent contigs reduced the number of contig gaps and more than doubled the N50 contig length as well as increased the number single-copy BUSCO genes (Table 1). With a more accurate representation of the haploid genome, the total length of the reference decreased by 55 Mbp (Table 1). It is noticeable that, with the exception of one linkage map produced here, these improvements were gained with a more efficient use of data generated for the original assembly. In addition to automated improvements with Lep-Anchor, we assembled and incorporated the native mitochondrial genome and used additional data from related individuals to characterize and reassemble a large inversion in LG19.

The improved assembly brings noticeable gains, and we could now successfully map the centromere associated repeat and unambiguously identify the centromere positions in all linkage groups (previously missing from LG1 and LG16, and incoherent in LG10 and LG14, Suppl. Fig. 2; see Varadharajan et al. (2019)). Removal of haplotypes and other changes in the genome assembly affects read mapping and single nucleotide variant calling. More even read depth and the anticipated mean depth indicate that the reference has become more haploid and contains fewer haplotype copies (Suppl. Fig. 3). In comparison to the ver. 6, the heterozygosity of the reference individual increased by 14% (Table 2), illustrating how the variation is concentrated in few regions and how these variable regions then get assembled as separate haplotypes. Indeed, most (78%) of the newly identified SNPs were in regions where haplotype variants were removed from the reference and reads from variant alleles now map to the same copy of the genomic region (Fig. 1b-c, Table 2). New SNPs in other regions were a minority and their allelic depth deviated from the expected (Suppl. Fig. 4).

Content of the LG12 changed considerably from ver. 6 to ver. 7 as one of the homologous copies in X and Y chromosomes were removed (Fig. 1a). As a result, the sex chromosome part of LG12 is close to a haploid representation and few regions show zero read depth (Fig. 2a). This also increases the heterozygosity of the male reference individual (Fig. 2a), the newly identified SNPs arising from differences between X and Y chromosomes, while no increase is observed in females (Fig. 2a). Although homologous sequences are represented only once, the sex chromosome is still a mosaic of X and Y chromosomes and females show both homozygous variant and reference alleles (Fig. 2a). An HMM analysis confirmed the mosaicism and indicated the sex chromosome assembly to be 57% of X chromosome (Suppl. Fig. 5). Despite the mosaicism, the reassembly improved the synteny of LG12 with the three-spined stickleback counterparts (Fig. 2b).

Scaffolding with Lep-Anchor had a minor impact on the variant allele frequencies in the LG19 inversion region (Fig. 3a). The reason for this is that the original contigs were mosaics of the two alleles and an improved ordering of contigs does not correct for their internal errors. The newly assembled contigs and the scaffolded alleles for the LG19 block revealed that there, indeed, are two segregating inversion haplotypes in the study population, and that the reference individual (see methods) is heterozygous (Fig. 3a). As expected, the variant allele frequencies across the newly assembled haplotypes are either zero or twice as high as with the original mosaic assembly for individuals homozygous for the two alleles (Fig. 3a). Although the mosaicism had a large impact on variant allele frequencies, its effect on SNP frequency was small. There are more SNPs according to ver. 7 but most of them are found due to haplotype removal and few of them are in the inversion region. With the HMM and data from the four females, we found four observable large regions that indicate fine-scale mosaic of two diverged haplotypes (Fig. 3b, see also Suppl. Table 3.). All these four regions were identified in both genome versions which suggests that the corresponding contigs are erroneously assembled as in LG19.

**FIGURE 3.**
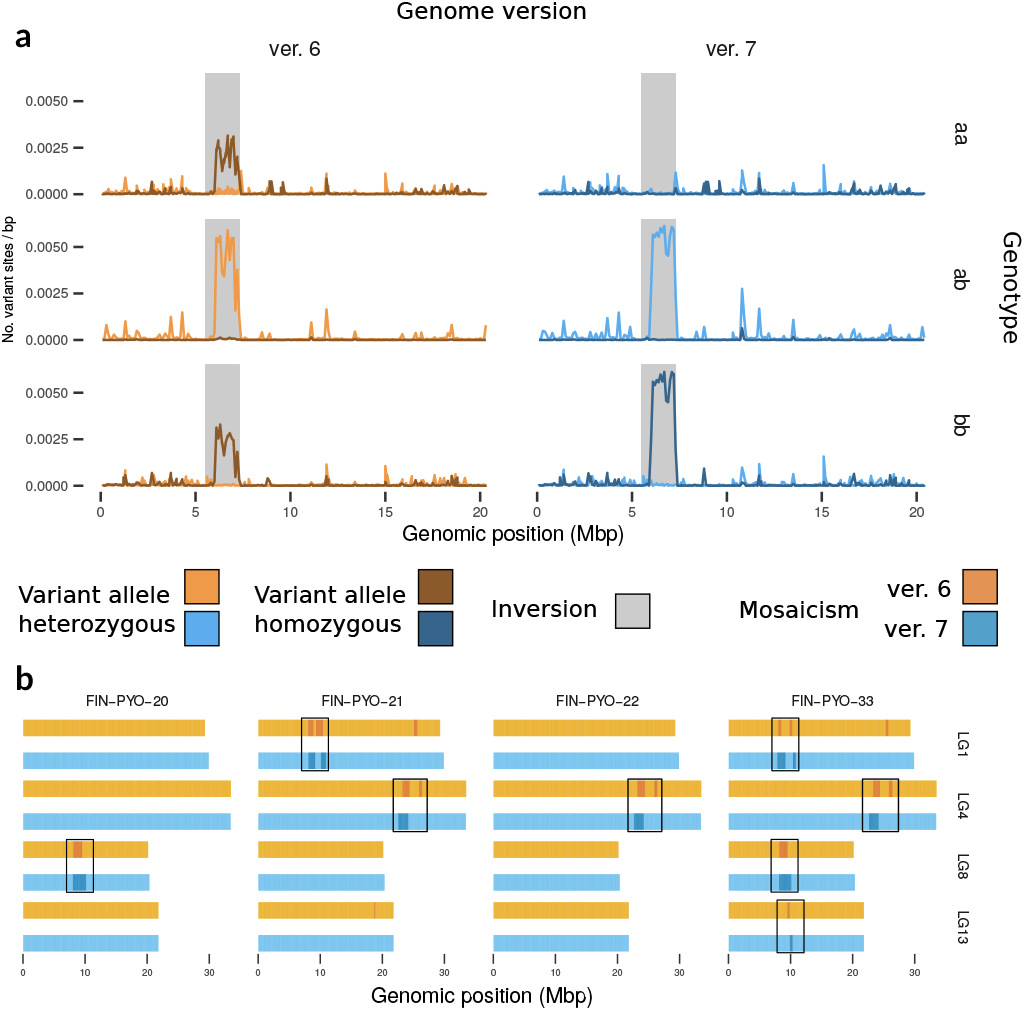
Examples of finescale mosaicism in the nine-spined stickleback reference geome. (a) When mapped to ver. 6, individuals homozygote for the LG19 inversion haplotypes (aa and bb; top and bottom) show high frequency of variant alleles in the inversion region. In ver. 7 with the reassembled inversion haplotype, individuals homozygous for the reference haplotype (top) have no variant alleles whereas those homozygous for the alternative allele (bottom) show two-fold frequency across the region. (b) Using a HMM, four candidates of finescale mosaicism (dark bands) similar to the LG19 inversion were identified. Here, the Viterbi path of the HMM algorithm is shown and only regions detected in both genome versions are highlighted with rectangles (see Suppl. Table 4 and Suppl. Fig. 6 for the genomic coordinates and posterior likelihoods).

Comparable data were not available for three-spined stickleback. We constructed a contig assembly by partitioning the full-length sequence (Peichel et al. 2017) at long runs of N’s, and constructed a linkage map for a distantly related population (Pritchard et al. 2017). In the absence of long-read data, we bridged the contigs of the original assembly using scaffolds of a 10X Genomics assembly (Berner et al. 2019). In this reassembly, we identified 1,831 haplotype contigs, most of them unassigned, and were able to add 176 previously unassigned contigs to the linkage groups. The ungapped length of the 21 linkage groups, representing the 21 chromosomes, decreased from 426 Mbp to 423 Mbp and the ungapped length of the unassigned contigs decreased from 21 Mbp to 13 Mbp. N50 of the original and our new genome are 83,717 and 87,370 bp, respectively (Suppl. Table 2). With the new reference we found 0.62% more autosomal SNPs in a sample from Paxton Lake, Canada, than were found using the original assembly (Table 2). Although the background heterozygosity of this individual was orders of magnitude higher than in our nine-spined stickleback reference individual, most of the newly identified SNPs (52%) were in regions where haplotype variant was removed from the reference genome. Whereas the median sequencing depth of the sample was 15x for both genome versions, the depth for the identified haplotype regions was 9x and 15x in the published and new assembly, respectively, indicating successful haplotype removal.

## DISCUSSION

Reconstructing a linear reference genome is a challenging, yet an instrumental task. Interpretation of genomic data is often made with the assumption that the reference genome is a complete haploid representation of the actual genome. The errors in the genome directly affect the conclusions drawn, and for instance, missing SNPs influence the site frequency spectrum that is essential in demographic analyses (Han, Sinsheimer, & Novembre 2014). More directly, presence of haplotype copies in a reference genome can make a highly diverged region seem exceptionally conserved and can thus seriously mislead variation-based functional analyses. Given the severe consequences of the errors, efforts to improve reference genomes are needed, and here we have described an approach to make reference genomes more haploid and more contiguous using the Lep-Anchor software (Rastas 2020).

Faced with the dilemma of correctly separating duplicated genome regions while simultaneously collapsing and merging haplotypic differences into a haploid sequence, all assembly programs are poised to make errors. The magnitude of these errors depends on the heterozygosity of the reference individual and on the type of input data, long reads spanning more distant sites and thus capable of creating longer haplotype blocks, while the direction of the bias to either too long or too short genome depends on the algorithm. While the three-spined stickleback genome is based on relatively old data and is established over years of refinement, the nine-spined stickleback genome is an example of a modern reference genome built using the best practices. We demonstrated our method’s potential by showing how the latter, an already very high quality reference genome, could be greatly improved by more efficient use of the original sequencing and mapping data (Fig.1, Table 1). Improvements were based on linking, reassembly and improved scaffolding of the contigs with joint use of linkage map anchoring and long read sequencing data, as well as characterization and removal of alternative haplotypes. The improvements on the three-spined stickleback genome were more modest but we could still both add new contigs into the linkage groups and remove haplotype copies (Suppl. Table 2, Suppl. Fig. 7), resulting in an 0.62% increase in number of segregating sites in a sample from the Paxton Lake benthic population (Table 2). We anticipate that the more modest changes in comparison to the nine-spined were due to absence of long reads and lower number of linkage map markers per contig in the three-spined data: 4.2 and 1.7 markers on average per contig in the nine and three-spined stickleback, respectively. While the three-spined stickleback analyses demonstrate that Lep-Anchor can improve even highly polished assemblies, they also illustrate how various data types, for example contigs from the 10X Genomics platform, can be incorporated in genome refinement.

In the nine-spined stickleback, most of the removed haplotypes were among the unassigned contigs and only one contig was moved between two linkage groups (Fig. 1a), underlining the high quality of the original scaffold. Although we were not able to place all contigs in the linkage groups, we were able to divide them in putative classes based on the read depth and their repeat content, those with high repeat content (either centromere or other) forming the largest groups of unassigned contigs. Although repetitive regions are difficult to assemble and scaffold using the type of data available, we were able to improve the centromeric regions (Fig. 1b) and our approach can be useful for repetitive regions more generally. Some unassigned contigs had low or even zero read depth, but as we did not detect any obvious contamination when aligning them to the NCBI database, those were retained in the reference genome.

Removal of haplotypes lead to identification of ca. 14% more autosomal SNPs in the nine-spined stickleback reference individual (Table 2). Finding more SNPs *per se* is not evidence for better assembly, and removal of true paralogous regions could lead to incorrect increase in SNP numbers. However, together with more uniform sequencing depth (Suppl. Fig. 3), strong evidence of successfully identified haplotypes (Fig. 1b-c) and higher number of single-copy BUSCOs (and lower number of duplicated BUSCOs, Table 1), our results show that genetic variability can be underestimated if the reference genome contains haplotypes. One should note, though, that our reference comes from a very small population and has extremely low background heterozygosity. Haplotypes, by definition, require variation between the copies and in our reference individual an exceptionally large proportion of the variation is concentrated within a small number of regions. The three-spined stickleback individual studied here had two orders of magnitude higher heterozygosity and, although the absolute numbers were larger, the relative impact of the reassembly on the SNP numbers was much smaller (Table 2). The minority of newly identified SNPs that were not within haplotype regions (22% of the novel SNPs in the nine-spined stickleback) may have emerged because of short similarities between contigs that were not classified as haplotypes. They may also be related to changes in mapping of the read pairs in regions where haplotype copies have been removed or contig orientation or order has changed. Nonetheless, the evidence for some of those SNPs is questionable as their allelic depth deviates from the expected (i.e. 0.5; Suppl. Fig. 4) and one may want to filter them from downstream analyses.

Nine-spined stickleback LG12 is formed by fusion of chromosomal segments that correspond to chromosomes 7 and 12 of the three-spined stickleback (Fig. 2b; Shikano et al. 2013). This rearrangement has occurred after the split of the three-spined and the nine-spined sticklebacks 17 million years ago (Guo et al. 2019) but the exact timing is unclear (Shikano et al. 2013). While 15 Mbp in one end of LG12 behaves like an autosomal chromosome, the 17 Mbp (25 Mbp in ver. 6) in the other end contains the sex-determination region and behaves like a sex chromosome (Fig. 2a). While parts of the sex-chromosome region seem very similar, other parts have differentiated significantly, and assembling complete X and Y chromosomes based on a single male reference individual is extremely challenging. Although our HMM analysis indicated that the LG12 assembled here is only 57% of X (Suppl. Fig. 5), we are confident that the sequence content of the current version is close to haploid presentation of X and the error is mainly in the SNP polarization. This is supported by the improved synteny with the three-spined stickleback genome (Fig. 2b) but especially by the more uniform read depth and more constant nucleotide diversity across the whole LG12 in females (Fig. 2a). The original sequencing data for the nine-spined stickleback reference are slightly outdated by modern standards, and we did not attempt to scaffold both X and Y copies of LG12. Fully separating the two should be relatively straightforward by obtaining long-read or linked-read data for both sexes with the latest sequencing technology.

Without genotype phasing, a haploid reference genome is a mosaic of maternal and paternal haplotypes and the reference alleles are drawn randomly. If parental haplotypes are clearly different, they are assembled as separate copies and appear as duplicates in the contig assembly; if the differences are punctuated by local similarities, the haploid consensus may alternate between the two parental haplotypes. It is evident that if the underlying contigs are erroneously assembled, their re-ordering cannot make the reference perfect. In the nine-spined stickleback, the inversion in LG19 and the sex-chromosome region LG12 demonstrate how diverged haplotypes complicate the assembly of a haploid reference genome. On the other hand, the characterization of the inversion haplotypes provides an example of how TrioBinning (Koren et al. 2018) can be utilized without a trio and long-read sequencing data combined with population level whole-genome sequencing data allow assembling the segregating haplotypes. We acknowledge that our HMM for identifying regions of diverged haplotypes provides only indicative results (Fig. 3b) but it does suggest that haplotypes can be fairly common in the nine-spined stickleback which is in line with findings regarding other fish (Stemple 2013) and humans (Sudmant et al. 2015). We also anticipate that highly concentrated alternation between two homozygous genotypes is a usable statistic for exploration and more sophisticated detection methods based on that could be devised. Identification of such regions requires the studied individual to be heterozygous and therefore all regions were not supported by all individuals. Having a single continuous haplotype, such as the inversion in LG19, in the reference genome correctly phases the alternative alleles (Fig. 3a) and allows studying the differences between the haplotypes. However, representation of potential structural differences is difficult and it is evident that methodological work to incorporate multiple haplotypes in a reference genome, e.g. using variation graph data structures, is urgently needed (Paten, Novak, Eizenga, & Garrison 2017).

Haploid reference genomes based on a single individual, such as the one here, represent only one version of the species’ genome which may cause reference bias and thus affect various downstream analyses and the conclusions drawn from them (Ballouz, Dobin, & Gillis 2019; Paten et al. 2017). Although a linear reference does not represent the full species diversity, they are widely used and provide a starting point for analysis of genomic variation between individuals and populations. In the future, pan-genome representations and graph-based algorithms will likely change the way reference genomes are represented and analyzed (Paten et al. 2017; Sherman & Salzberg 2020). Since linear genomes are still widely used, their improvements are relevant and our work demonstrates that significant enhancements can be obtained with efficient use of the existing data. Moreover, characterization of haplotypes is instrumental in more inclusive genome representations, increasing the relevance of our approach.

## DATA AVAILABILITY

Sequence data that support the findings of this study have been deposited in European Nucleotide Archive (ENA) under the accessions PRJEB39736 (linkage map parents), PRJEB39760 (linkage map offspring), PRJEB33474 and PRJEB39599 (whole-genome sequenced individuals from Pyöreälampi and Kirkasvetinen). Linkage maps and genomes of the nine- and three-spined sticklebacks are available at https://github.com/mikkokivikoski/NSP_V7 and in ENA under the accession GCA_902500615. Computer code and instructions for automated improvement of reference genome assemblies and scripts for reproducing these analyses are available at https://github.com/mikkokivikoski/NSP_V7.

## ACKNOWLEDGEMENTS

We thank Chris Eberlein, Sami Karja, Takahito Shikano, and everyone else who helped in catching and rearing the fish utilized in this work. Thanks are also due to those who helped in DNA extractions, Laura Hänninen, Kirsi Kähkönen, Miinastiina Issakainen and Sami Karja in particular. We thank Antoine Fraimout for help with relatedness analysis. Our research was supported by LUOVA doctoral programme of the University of Helsinki (funding to MK), Academy of Finland (# 129662, 134728 and 218343 to JM; # 322681 to AL), and grant from Helsinki Institute for Life Sciences (HiLife; to JM). We also wish to acknowledge CSC – IT Center for Science, Finland, for computational resources.

## AUTHOR CONTRIBUTIONS

J.M. directed and organized the data collection. P.R. generated linkage maps and assembled reference genomes. M.K., A.L. and P.R. carried out the data analysis and interpretation of the results. M.K. was responsible for structuring and compiling the manuscript. The manuscript was written and edited by all authors.

